## Supplementary Text 1 for "Mechanistic model for human brain metabolism and its connection to the neurovascular coupling"

**Supplementary example to highlight the issues with parameter unidentifiability**

To illustrate the problem with unidentifiability brought up in the discussion section, we have created a two-compartment version of the model presented in this paper (S1 Fig). This two-compartment model is fitted to the experimental data for the metabolic response (section 2.5.1) and the parameter distributions are determined in the manner described in sections 2.4.1 and 2.4.2 in the manuscript. If we compare the estimated distributions of four parameters (${k_{max}}_{pyr2}, K_{M_{Pyr2}}, k_{max_{Glut1}}, k_{max_{Glut2}}$), it becomes apparent that the parameter distributions of the single compartment model (S2 Fig, blue) are more well determined than the corresponding parameter distributions of the two-compartment model (S2 Fig red and yellow). Supplementary figure 2 shows a comparison of these parameter distributions in the form of a box chart. In this figure, the distributions are plotted on the y-axis with the different parameters considered on the x-axis. The blue boxes represent the parameter distributions for the single compartment model, the red boxes show the distributions for the corresponding parameters in the neuron compartment of the two-compartment model, and the yellow boxes show the distributions for the corresponding parameters in the astrocyte compartment of the two -compartment model. As can be seen, the red and yellow distributions of the two-compartment model are several orders of magnitude wider than the corresponding bule distributions of the single compartment model (note that the y-axis is plotted in log-scale). This problem appears since the parameters in the original single compartment model represent a combination of the parameters in the two-compartment model. It should be noted that this problem of identifiability is the case for a number of parameters in the two-compartment model, including but not limited to the four parameters showcased in the example.

This example shows that adding additional complexity to the model in the form of multiple compartments i) does not improve the model’s ability to explain the data we have considered in the manuscript, but ii) that it does reduce the parameter identifiability. The same problem would likely be the case if we added other excluded aspects of the metabolism such as, glycogen metabolism, or expanded the metabolic pathways that are currently lumped together into single reaction.

Note that these results and remarks do not mean that these additional processes do not occur in the brain, or that we claim that they do not occur. There are several works that point to the importance of these processes for the cerebral metabolism. However, given the experimental data considered in this work, it is not possible to model these mechanisms and preserve well determined parameters. To achieve well determined parameters for a more comprehensive model we would need to consider additional data that could be used to limit the parameter solution space. This could of course be done, but there is a limit to how much published data is relevant to consider, and there is a clear limit on what is currently available, measured on real human subjects. We think that the amount of data we have presented herein is relevant for the current aims of this work. Having said this, considering additional data from other sources to expand the model is something that would be highly interesting in future works.

Finally, all models of biological systems are simplifications to some degree, and, in our modelling approach, the available experimental data is the factor that determines the degree of simplification. In other words, even if we would add the mechanisms such as glycogen metabolism, multiple compartments, etc, the resulting model would still neglect the majority of existing knowledge that exists about these systems (e.g. that there is a wide range of different cell types, many more than two; that there are many more reactions that happens that are still assumed negligible or not important to include as independent reactions, etc). Thus, these additions would add more knowledge, but would a) remove us from presenting a minimal model, with identifiable parameters, b) still be possible to expand in the same way to add further known mechanisms. We therefore think that it is more clearcut to present a minimal model, since this has the added benefit of identifiability of parameters.
