## Supplementary Table 1 for "Mechanistic model for human brain metabolism and its connection to the neurovascular coupling"

### S2 Table: Implementation of Sten *et al.* 2017 model

| **State Equations** | **Interpretation** |
| --- | --- |
| $\frac{d(Stimulus)}{dt} = 0$ | Stimulus input signal |
| $\frac{d(oHb)}{dt}= {v1}_{b}- {v1}_{f}+v_{inoHb}-v_{outoHb}$ | Change in oxyhemoglobin level |
| $\frac{d(dHb)}{dt}={v1}_{f}- {v1}_{b}+v_{indHb}-v_{outdHb}$ | Change in deoxyhemoglobin level |
| $\frac{d\left( O_{2A} \right)}{dt}=v_{O_{2A}} \times v_{bv}-v_{basalMet} \times k_{prop1}- v_{stimMet} \times k_{prop2}*k_{volumeScale2}$ | Change in free oxygen in blood compartment |
| $\frac{d\left( O_{2B} \right)}{dt}={v1}_{f}- {v1}_{b}-v_{O_{2A}}+v_{inO_{2}}-v_{outO_{2}}$ | Change in free oxygen in extravascular tissue |
| $\frac{d\left( {Glucose}_{A} \right)}{dt}={(v}_{Gluc1}+ v_{Gluc2})v_{bv}*k_{volumeScale}-v_{basalMet}- v_{stimMet}$ | Change in glucose (blood compartment) |
| $\frac{d\left( {Glucose}_{B} \right)}{dt}=-\left( v_{Gluc1}+ v_{Gluc2} \right)+v_{inG}-v_{outG}$ | Change in glucose level in extravascular tissue |
| $\frac{d\left( {Delay}_{M} \right)}{dt}= {input}_{1}-{met}_{sink}$ | Delay state |
| $\frac{d(Glutamate)}{dt} = {input}_{2} - {Glutamate}_{sink}$ | Glutamate release upon stimulation |
| $\frac{d(GABA)}{dt} = {input}_{3} - {GABA}_{sink}$ | GABA release upon stimulation |
| $\frac{d\left( {Ca}^{2+} \right)}{dt} = k_{Ca}\left( 1+v_{glutamate} \right)$  $\left( \frac{1}{1+v_{GABA}\left( 1+v_{diazepam} \right)} \right)-{Ca}_{sink}$ | Calcium influx in the astrocyte |
| $\frac{d(AA)}{dt} = v_{calcium}-\left( k_{vc}+k_{vd} \right)AA$ | Change in AA level |
| $\frac{d(\text{AA-met}_{\text{vc1}})}{dt} = k_{vc}AA-v_{vc1}$  $\frac{d(\text{AA-met}_{\text{vc2}})}{dt} = v_{vc1}-v_{vc2}$  $\frac{d(\text{AA-met}_{\text{vc3}})}{dt} = v_{vc2}-v_{vc3}$  $\frac{d(\text{AA-met}_{\text{vc4}})}{dt} = v_{vc3}-v_{vc4}$ | Intermediary states |
| $\frac{d(\text{AA-met}_{\text{vd1}})}{dt} = k_{vd}AA-v_{vd1}$  $\frac{d(\text{AA-met}_{\text{vd2}})}{dt} = v_{vd1}-v_{vd2}$  $\frac{d(\text{AA-met}_{\text{vd3}})}{dt} = v_{vd2}-v_{vd3}$  $\frac{d(\text{AA-met}_{\text{vd4}})}{dt} = v_{vd3}-v_{vd4}$ | Intermediary states |
| **Reactions** | **Interpretation** |
| ${v1}_{f} = {k1}_{f} \times oHb$  ${v1}_{b} = {k1}_{b} \times dHb \times O_{2B}$ | Rate of releasing oxyhemoglobin into oxygen and deoxyhemoglobin  Rate of binding oxygen and deoxyhemoglobin into oxyhemoglobin |
| $v_{inoHb} = {oHb}_{basal} \times v_{flow}$  $v_{outoHb} = oHb \times v_{flow}$  $v_{indHb} = {dHb}_{basal} \times v_{flow}$  $v_{outdHb} = dHb \times v_{flow}$  $v_{inG} = {{(Glucose}_{B})}_{basal} \times v_{flow}$  $v_{outG} = {Glucose}_{B} \times v_{flow}$  $v_{inO_{2}} = {{(O}_{2B})}_{basal} \times v_{flow}$  $v_{outO_{2}} = O_{2B} \times v_{flow}$ | Oxyhemoglobin influx  Oxyhemoglobin outflux  Deoxyhemoglobin influx  Deoxyhemoglobin outflux  Glucose influx  Glucose outflux  Oxygen influx  Oxygen outflux |
| $v_{O_{2A}}=k_{O2}\left( O_{2B}-O_{2A} \right)$  $v_{Gluc1}=k_{Gluc1}\left( {Glucose}_{B}-{Glucose}_{A} \right)$  $v_{Gluc2}=k_{Gluc2}\left( \frac{{Glucose}_{B}}{k_{m}+{Glucose}_{B}} \right)$ | Diffusion of oxygen between blood and cell compartment  Diffusion of glucose between blood and cell compartment  Receptor mediated transportation of glucose between blood and cell compartment |
| $v_{basalMet} = k_{basalMet} \times{O_{2A}}^{prop1} \times{Glucose}_{A}$  $v_{stimMet} = {Delay}_{M} \times{O_{2A}}^{prop2} \times{Glucose}_{A}$ | Basal metabolism  Stimulation induced metabolism |
| ${input}_{1} = k_{met} \times Stimulus$  ${input}_{2} = k_{Glutamate}\times Stimulus$  ${input}_{3} = k_{GABA}\times Stimulus$  ${Glutamate}_{Sink} = {sink}_{Glutamate} \times Glutamate$  ${GABA}_{Sink} = {sink}_{GABA} \times GABA$  ${met}_{Sink} ={sink}_{met} \times{Delay}_{M}$ | Stimulus input to the metabolic module  Stimulus input to glutamate release  Stimulus input to GABA release  Glutamate degradation  GABA degradation  Degradation of increased metabolism |
| $v_{glutamate} = k_{3} \times Glutamate$  $v_{GABA} = \frac{GABA}{k_{4}}$  $v_{diazepam} = \frac{c_{diaz}^{n}}{k_{diaz}^{n}+c_{diaz}^{n}}$  ${Ca}_{sink} = {sink}_{Ca} \times{Ca}^{2+}$  $v_{calcium}= k_{PL} \times{Ca}^{2+}$ | Glutamate’s effect on calcium  GABA’s effect on calcium  Diazepam’s effect on calcium  Degradation of calcium  Calcium’s effect on AA |
| $v_{vc1} = k_{vc1} \times\text{AA-met}_{\text{vc1}}$  $v_{vc2} = k_{vc2} \times\text{ AA-met}_{\text{vc2}}$  $v_{vc3} = k_{vc3} \times\text{ AA-met}_{\text{vc3}}$  $v_{vc4} =k_{vc4} \times\text{ AA-met}_{\text{vc4}}$ | Delay states |
| $v_{vd1} = k_{vd1} \times\text{AA-met}_{\text{vd1}}$  $v_{vd2} = k_{vd2} \times\text{ AA-met}_{\text{vd2}}$  $v_{vd3} = k_{vd3} \times\text{ AA-met}_{\text{vd3}}$  $v_{vd4} =k_{vd4} \times\text{ AA-met}_{\text{vd4}}$ | Delay states |
| **Variable values** | **Interpretation** |
| ${v_{flow}=k}_{flow} + e^{k_{s} \times\text{AA-met}_{\text{vd4}} - {k_{i} \times\text{AA-met}}_{\text{vc4}}}$ | Blood flow |
| ${v_{bv}=\left( v_{flow} \right)}^{k_{bv}}$ | Blood volume |
| $Glucos{e_{B}}_{basal}=$10 | Basal amount of glucose in blood |
| $oHb_{basal}=$10 | Basal amount of oxygen in blood |
| ${O_{2B}}_{basal}=$10 | Basal amount of oxygenated hemoglobin in blood |
| $dHb_{basal}=$10 | Basal amount of deoxygenated hemoglobin in blood |
| $BOLD=e^{-k_{y} \times dHb}$ | Measurement signal |
