## Supplementary Table 2 for "Mechanistic model for human brain metabolism and its connection to the neurovascular coupling"

### S2 Table: Parameter values

| Parameter Name | Unit | Optimal Parameter Value | Upper parameter bound | Lower parameter bound |
| --- | --- | --- | --- | --- |
| ky | unitless | 7.3000 | 56.820 | 0.0100 |
| kmetabolic | 1/s | 0.0002 | 0.3200 | 0.0001 |
| k1f | 1/(amount$\times$s) | 0.0003 | 0.0810 | 0.0001 |
| k1b | 1/(amount$\times$s) | 0.1300 | 3.1100 | 0.0010 |
| $k_{basalMet}$ | 1/s | 0.0500 | 0.0070 | 0.0010 |
| kflowBasal | amount/s | 0.0600 | 0.8900 | 0.0001 |
| $k_{prop1}$ | Ratio of oxygen/glucose metabolised | 0.1200 | 1.1000 | 0.0001 |
| $k_{prop2}$ | Ratio of oxygen/glucose metabolized | 0.0100 | 0.3500 | 0.0001 |
| kO2bbb | 1/s | 0.0600 | 20.670 | 0.0001 |
| kGlucf | 1/s | 0.9000 | 22.760 | 0.0020 |
| kGlucb | 1/s | 0.0001 | 0.0100 | 0.0001 |
| kBV | unitless | 1.6000 | 96.820 | 0.0001 |
| Kmgluc | amount | 0.0100 | 10.240 | 0.0001 |
| kGD | 1/s | 0.0600 | 95.270 | 0.0200 |
| PL | 1/s | 0.1200 | 69911.0 | 0.0001 |
| CaBas | 1/s | 0.0200 | 32615.0 | 0.0001 |
| sinkGABA | 1/s | 0.2500 | 2.3200 | 0.0001 |
| sinkGlu | 1/s | 1.0200 | 5.3900 | 0.2400 |
| sinkA | 1/s | 1.2300 | 7.2300 | 0.3400 |
| sinkCon | 1/s | 0.8800 | 9.5100 | 0.3000 |
| sinkDil | 1/s | 0.6400 | 69990.0 | 0.2400 |
| Kglut | 1/s | 0.1300 | 0.3200 | 0.0010 |
| Kgaba | 1/s | 0.0001 | 0.4700 | 0.0001 |
| k3 | 1/s | 886.93 | 69999.0 | 427.37 |
| k4 | amount$\times$s | 26.710 | 26222.0 | 0.0002 |
| k5 | 1/s | 1.0100 | 7.3000 | 0.2800 |
| k7 | 1/s | 0.1100 | 1.0500 | 0.0001 |
| b3 | unitless | 1.1800 | 69994.0 | 0.0100 |
| b4 | unitless | 0.2200 | 214.41 | 0.0001 |
| kdelay1d | 1/s | 0.4400 | 4287.7 | 0.1400 |
| kdelay2d | 1/s | 55521.0 | 69999.0 | 0.5400 |
| kdelay3d | 1/s | 2.5400 | 59.820 | 0.1300 |
| kdelay1c | 1/s | 1.2100 | 7.2800 | 0.3600 |
| kdelay2c | 1/s | 0.9100 | 10.760 | 0.3900 |
| kdelay3c | 1/s | 1.4900 | 108.09 | 0.4600 |
| ${k_{max}}_{pyr}$ | 1/s | 61.230 | 7069.9 | 0.0100 |
| ${K_{M}}_{pyr}$ | amount | 0.7700 | 69266.0 | 0.0030 |
| k1 | 1/s | 0.0050 | 0.0100 | 0.0020 |
| ky1 | Unitless | 0.0001 | 42201.0 | 0.0001 |
| ky2 | Unitless | 69890.0 | 69999.0 | 0.0800 |
| ${k_{max}}_{OAA}$ | 1/s | 0.0007 | 0.0100 | 0.0001 |
| ${K_{M}}_{OAA}$ | amount | 0.0001 | 0.1500 | 0.0001 |
| ${k_{max}}_{OG1}$ | 1/s | 0.1900 | 69964.0 | 0.0100 |
| ${k_{max}}_{OG2}$ | 1/s | 0.0200 | 31384.0 | 0.0004 |
| ${k_{max}}_{Glut1}$ | 1/s | 0.0300 | 2.9100 | 0.0001 |
| ${k_{max}}_{Glut2}$ | 1/s | 0.0010 | 0.0040 | 0.0001 |
| ${k_{max}}_{Gln}$ | 1/s | 0.0200 | 2.1800 | 0.0001 |
| ${k_{max}}_{Asp}$ | 1/s | 0.0300 | 69900.0 | 0.0004 |
| ${k_{max}}_{PO}$ | 1/s | 14.290 | 69980.0 | 0.0100 |
| ky3 | Unitless | 1993.3 | 69854.0 | 0.0001 |
| ky4 | Unitless | 46.830 | 69990.0 | 0.0100 |
| ${k_{max}}_{Pyr2}$ | 1/s | 0.0200 | 0.2600 | 0.0001 |
| $k_{stim1}$ | Unitless | 0.0060 | 69981.0 | 0.0001 |
| $k_{stim2}$ | Unitless | 2588.0 | 69830.0 | 0.0001 |
| ${K_{M}}_{Pyr2}$ | Amount | 0.0020 | 0.4400 | 0.0001 |
| k_volumeScale | Unitless | 0.4900 | 8.5900 | 0.0002 |
| k_volumeScale2 | Unitless | 5.8600 | 788.74 | 0.0001 |
| Neg_kglut | 1/s | 0.1800 | 55584.0 | 0.0010 |
| Neg_kgaba | 1/s | 714.54 | 69999.0 | 0.0400 |
| Neg_kmetabolic | 1/s | 0.0010 | 69788.0 | 0.0001 |
| Neg_kdelay2d | 1/s | 0.3000 | 406.71 | 0.0900 |
| Neg_kdelay2c | 1/s | 0.0500 | 0.1400 | 0.0001 |
