## Supplementary Zip file for "Mechanistic model for human brain metabolism and its connection to the neurovascular coupling": GenerateData.m

```
%Generate Data-structure used in simulation
function [metData,BOLD_data,bednData,grutData] = GenerateData()
addpath(genpath(fullfile(pwd,'Data')));

%% get metabolite data
load('CalcSEdoneDoubleData.mat');
metData(1).t=round(linDoubleData.Time*60);
metData(1).Y=[linDoubleData.LacMean linDoubleData.GluMean linDoubleData.GlcMean linDoubleData.AspMean zeros(length(metData(1).t),1)];
metData(1).Sigma_Y=[linDoubleData.LacSE linDoubleData.GluSE linDoubleData.GlcSE linDoubleData.AspSE inf(length(metData(1).t),1)];

load('CalcSEdoneSingleData.mat');
metData(2).t=round(linSingleData.Time*60);
metData(2).Y=[linSingleData.LacMean linSingleData.GluMean linSingleData.GlcMean linSingleData.AspMean zeros(length(metData(2).t),1)];
metData(2).Sigma_Y=[linSingleData.LacSE linSingleData.GluSE linSingleData.GlcSE linSingleData.AspSE inf(length(metData(2).t),1)];

%% get fMRI data
load('fastfMRI_BOLDdata.mat');
BOLD_data(1).t = 0:0.7:((28-1)*0.7)';
BOLD_data(1).Y = [zeros(28,4), fastfMRI_BOLDdata.visMean'];
BOLD_data(1).Sigma_Y = [inf(28,4), [1;fastfMRI_BOLDdata.visSE(2:end)']];

BOLD_data(2).t = 0:0.7:((28-1)*0.7)';
BOLD_data(2).Y = [zeros(28,4), fastfMRI_BOLDdata.dmnMean'];
BOLD_data(2).Sigma_Y = [inf(28,4), [1;fastfMRI_BOLDdata.dmnSE(2:end)']];


%% get bednarik Data
load('bednarikData.mat');
bednData(1).t=round(bednarikData.Time*60);
bednData(1).Y=[((bednarikData.LacMean/mean(bednarikData.LacMean(1:2,1))-1)*100) ((bednarikData.GluMean/mean(bednarikData.GluMean(1:2,1))-1)*100) ((bednarikData.GlcMean/mean(bednarikData.GlcMean(1:2,1))-1)*100) ((bednarikData.AspMean/mean(bednarikData.AspMean(1:2,1))-1)*100)];
bednData(1).Sigma_Y=[((bednarikData.LacSE/(mean(bednarikData.LacMean(1:2,1))))*100) ((bednarikData.GluSE/(mean(bednarikData.GluMean(1:2,1))))*100) ((bednarikData.GlcSE/(mean(bednarikData.GlcMean(1:2,1))))*100) ((bednarikData.AspSE/(mean(bednarikData.AspMean(1:2,1))))*100)];


%% get grutter data
load('GruetterData.mat');
grutData(1).t=round(GruetterData.Time*60);
grutData(1).Y=[((GruetterData.LacMean/(sum(GruetterData.LacMean(1:15,1))/15)-1)*100) ((GruetterData.GluMean/(sum(GruetterData.GluMean(1:15,1))/15)-1)*100)];
grutData(1).Sigma_Y=[((GruetterData.LacSE/((sum(GruetterData.LacMean(1:15,1)))/15))*100) ((GruetterData.GluSE/((sum(GruetterData.GluMean(1:15,1)))/15))*100)];

end
```
