## Supplementary Zip file for "Mechanistic model for human brain metabolism and its connection to the neurovascular coupling": GenerateModels.m

```
function []=GenerateModels(modelName)
% run('../AMICI-0.10.11_SS_eventFix/matlab/installAMICI.m')% set path for the toolbox.
basePath =pwd;
cd(sprintf('./Models/%s',modelName))
symsFiles = dir('*_syms.m');% find which _syms.m files exists.
symsFileNames = vertcat({symsFiles.name})';
modelNames = cellfun(@strrep,symsFileNames,repmat({'_syms.m'},size(symsFileNames)),repmat({''},size(symsFileNames)),'uni',0); % get the model name without the _syms.m surfix

%% find which model are already exists as .mex* files 

mexFiles = dir('*.mex*'); % find which mex files exists.
mexFileNames = vertcat({mexFiles.name})';

mexFileExistsIndx = false(1,size(modelNames,1));
mexIndx = cellfun(@strfind,mexFileNames,repmat({'.mex'},size(mexFileNames)))-1; % find the index of the .mex* surfix

for i = 1:length(mexFileNames)%find the logical idexing for the models that already have existing .mex* files
mexFileExistsIndx(strcmp(mexFileNames{i}(5:mexIndx(i)),modelNames))=true;
end

%% Compile the models that don't have an existing .mex* file  

for i = find(~mexFileExistsIndx)
   fprintf('\n\nCompiling model: %s\n',modelNames{i});
   try
       amiwrap(modelNames{i},symsFileNames{i}(1:end-2),pwd)
       fprintf('\nModel %s sucessfully compiled.\n',modelNames{i});
   catch ErrorReport
       disp(getReport(ErrorReport));
       fprintf('\nModel %s compilation failed!.\n',modelNames{i}); 
   end   
end

cd(basePath);
end 
```
