## Supplementary Zip file for "Mechanistic model for human brain metabolism and its connection to the neurovascular coupling": maximizeAxes.m

```
function [Figure] = maximizeAxes(Figure,n,m,axOrder)
% [Figure] = untitled(Figure,n,m) - adjusts the outerpositions of the axes
% in the figure such that the figure margin is minimised. 
% Figure is a matlab figure handel
% n and m are the number of rows and columnd of subplots presens. 
% axOrder is an index matrix which is used if the order of the subplots
% differ from the matlab default order.

if ~isa(Figure,'matlab.ui.Figure')
error('%s is NOT a valid figure handle!',inputname(1));
return
else
   axesIndex = false(length(Figure.Children),1);
   for k = 1:length(Figure.Children)
       axesIndex(k) = isa(Figure.Children(k),'matlab.graphics.axis.Axes');
   end
   axes = flipud(Figure.Children(axesIndex));
   
   if  length(axes) > n*m
       error('inputs n=%d and m=%d does not match the number of axes in figure %s',n,m,inputname(1));
       error('Make sure that %s corresopnds to the number of rows of subplots and %s corresopnds to the number of columns of subplots',inputname(2),inputname(3))
       return
   else
       if nargin < 4 || isempty(axOrder)
           axOrder = [1:length(axes)];
       elseif nargin == 4 && ~isempty(axOrder)
           if length(unique(axOrder)) ~= length(axes)
               error('There are %d axes in the figure but only %d unique indecies in the vector axOrder! Ensure that the two match in length and that all elements in axOrder are uique.',length(axes),length(unique(axOrder)));
               return
           end
       end
       if length(axes) < n*m
           for i = (n*m)-length(axes):-1:1
              subplot(n,m,(m*n+1)-1)
          end
       end
       
       AxPositions=cell(n*m,1);
       for i = 1:n
           for j = 0:(m-1)
               AxPositions(i+j+(i-1)*(m-1))={[(j*1/m),((n-i)/n),(1/m),(1/n)]};
           end
       end
       set(axes,{'OuterPosition'},AxPositions(axOrder));
   end
   
   %Aling the x.positions and the widths of the columns
   for i = 1:m
       for j = 0:n-1
           axes(i+j*m).Position([1,3])= axes(i).Position([1,3]);
       end
   end
   
   %Aling the y.positions and the hights of the rows
   for i = 1:n
       for j = (1:m)+(i-1)*m
           axes(j).Position([2,4])= axes(1+m*(i-1)).Position([2,4]);
       end
   end
   
end

end
```
