## Supplementary Zip file for "Mechanistic model for human brain metabolism and its connection to the neurovascular coupling": setSubFigureNotations.m

```
function [stHandel]=setSubFigureNotations(ax,figLabelFontsize)
    %% add the A-F notations to the topleft corner of axes
    % figLabelFontsize = 30;
    sgTitles = cellfun(@char,num2cell(65:65+length(ax)-1),'uni',false);
    for i = 1:length(ax)
        stHandel(i) = annotation('textbox',[.01 .1 .3 .3],'String',sgTitles{i},'FitBoxToText','on','FontSize',figLabelFontsize,'FontWeight','bold','LineStyle','none');
        stHandel(i).Position(1:2)=[ax(i).Position(1),ax(i).Position(2)+ax(i).Position(4)-stHandel(i).Position(4)];
    end
end
```
