## Supplementary Zip file for "Mechanistic model for human brain metabolism and its connection to the neurovascular coupling": MCMC_sampling.m

```
%% set Paths
clear all 
close all

% set path for the toolbox.
addpath(genpath(fullfile(pwd,'lib')))
addpath(genpath(fullfile(pwd,'Models')))
addpath(genpath(fullfile(pwd)))

modelName = 'NVC_Metabolism_model_V6';
%% load start parametres
% load('./Results/20200930-202957/NVC_Metabolism_model_V6/opt(213.44)_201001-034806.mat')
load('./Parameters/opt(213.44)_201001-034806_v6.mat')
theta = optParamM;

%% load Data
[metData,BOLD_data,~,~] = GenerateData(); % Get the metabolite and BOLD data
load('.\Data\lundengardData.mat')
%% Here i've in a previous article specified sim options outside of costfunction, do as you wish
% set amiOptions
options=amioption();
options.maxsteps=1e4;
options.atol=1e-16;
options.rtol=1e-8;

%% Create file where all chi-2 acceptable parameters are saved
FileName=sprintf('MCMC_%s_%s.dat',modelName,datestr(now,'yymmdd_HHMMSS')); %use for real run.
% FileName = 'TempTESTFile.dat'; % use for testing and debugging such that a new file is not created for each run.
FID = fopen(FileName,'wt');


%% Set the objective function
objectiveFunction=@(X) MetCost(X,metData,BOLD_data,modelName,options,0,FID);


%% MCMC sampling using the rampart algorithm
% One of the more advanced algorithms but it is nicely implemented in the
% PESTO toolbox. The following script starts the monte
% carlo sampling using the best identified parameter vector (which you load) as starting
% point, All results are saved in the parameters structure, additionally, a file is created and stores all chi-2 acceptable parameter vectors, with their associated cost.
%% Initial setup
% the toolbox PESTO uses this parameters structure, list all parameters names for automatic plottning
for i=1:length(theta) %For every parameter
parameters.name{i} = num2str(i);
end
%% Set parameter bounds 

parameters.min = log10(1e-4*ones(size(theta)));
parameters.max = log10(7e4*ones(size(theta)));

parameters.number = length(parameters.name);           % number of parameters

%% Options
optionsPesto = PestoOptions(); % loads optimization options and options for everything basically

optionsPesto.obj_type = 'negative log-posterior';  % Works with least squares cost which we typically been using, i.e the goal is the minimize. so keep it as is
optionsPesto.n_starts = 1; % number of multi-starts optimization performed <-- ignore this, better to start sampling in a previously estimated optima
optionsPesto.mode = 'visual'; %'silent'' it will plot the results itteratively. 

optionsPesto.comp_type = 'sequential'; % no parallel computing 
%% If using SBToolbox, it won't return everything as needed (i.e hessian and derivatives, have this line active)
% The algorithm does not need any sensitivities to work, therefore only one
% output
optionsPesto.objOutNumber=1;

%% Markov Chain Monte Carlo sampling -- Parameters
% Values for the parameters are sampled by using an advanced Parallel Tempering (PT)
% algorithm. This way, the underlying probability density of the parameter 
% distribution can be captured. if only ONE temperature is used, this is
% effectively an adapted Metropolis algorithm single-chain algorithm.

% Building a struct covering all sampling options:
optionsPesto.MCMC = PestoSamplingOptions();
optionsPesto.MCMC.nIterations = 1e6; %% This is the most important setting, covering how many iterations to be performed, here i think the minimum is 1e5. Gold standard is usually 1e6 iterations.
optionsPesto.MCMC.mode = optionsPesto.mode;
%% Using RAMPART  Most of this should be left as is, however look in the file called RAMPARTOPTIONS in @RAMPARTOPTIONS folder to see what each setting does
      optionsPesto.MCMC.samplingAlgorithm     = 'RAMPART';
      optionsPesto.MCMC.RAMPART.nTemps           = 62;
      optionsPesto.MCMC.RAMPART.exponentT        = 1000;
      optionsPesto.MCMC.RAMPART.maxT             = 2000;
      optionsPesto.MCMC.RAMPART.alpha            = 0.51;
      optionsPesto.MCMC.RAMPART.temperatureNu    = 1e3;
      optionsPesto.MCMC.RAMPART.memoryLength     = 1;
      optionsPesto.MCMC.RAMPART.regFactor        = 1e-8;
      optionsPesto.MCMC.RAMPART.temperatureEta   = 10;
      
      optionsPesto.MCMC.RAMPART.trainPhaseFrac   = 0.1;
      optionsPesto.MCMC.RAMPART.nTrainReplicates = 5;
      
      optionsPesto.MCMC.RAMPART.RPOpt.rng                  = 1;
      optionsPesto.MCMC.RAMPART.RPOpt.nSample              = floor(optionsPesto.MCMC.nIterations*optionsPesto.MCMC.RAMPART.trainPhaseFrac)-1;
      optionsPesto.MCMC.RAMPART.RPOpt.crossValFraction     = 0.2;
      optionsPesto.MCMC.RAMPART.RPOpt.modeNumberCandidates = 1:20;
      optionsPesto.MCMC.RAMPART.RPOpt.displayMode          = 'text';
      optionsPesto.MCMC.RAMPART.RPOpt.maxEMiterations      = 100;
      optionsPesto.MCMC.RAMPART.RPOpt.nDim                 = parameters.number;
      optionsPesto.MCMC.RAMPART.RPOpt.nSubsetSize          = 1000;
      optionsPesto.MCMC.RAMPART.RPOpt.lowerBound           = parameters.min;
      optionsPesto.MCMC.RAMPART.RPOpt.upperBound           = parameters.max;
      optionsPesto.MCMC.RAMPART.RPOpt.tolMu                = 1e-4 * (parameters.max(1)-parameters.min(1));
      optionsPesto.MCMC.RAMPART.RPOpt.tolSigma             = 1e-2 * (parameters.max(1)-parameters.min(1));
      optionsPesto.MCMC.RAMPART.RPOpt.dimensionsToPlot     = [1,2];
      optionsPesto.MCMC.RAMPART.RPOpt.isInformative        = [1,1,ones(1,optionsPesto.MCMC.RAMPART.RPOpt.nDim-2)];
      


      %% Here important part, these two lines are only valid if you have done the multi-start opt
%optionsPesto.MCMC.theta0 = parameters.MS.par(:,1); 
%optionsPesto.MCMC.sigma0 = 0.5 * inv(squeeze(parameters.MS.hessian(:,:,1)));

%% if multi-start does not work / is not used,  use these two lines
 optionsPesto.MCMC.theta0 = theta;
optionsPesto.MCMC.sigma0 = 1e5 * eye(length(theta));


%% Run the sampling
warning('off','all')
parameters = getParameterSamples(parameters, objectiveFunction, optionsPesto);
warning('on','all')

%% Create folder with name MCMC_DAY-MONTH-YEAR hour-min-sec (Very advanced matlab)
% fclose(FID);
folderStr=datestr(now,'yymmdd_HHMMSS');
folderStr=strrep(folderStr,{':',' '},'_');
folderStr=folderStr{2};
mkdir(['./Results/MCMCResult_' folderStr])
FolderName=fullfile(['./Results/MCMCResult_' folderStr]);
%% Save results to folder
save(fullfile(FolderName,'parameters.mat'),'parameters')
fclose(FID);
movefile(FileName,FolderName);
%% Takes all open figures and saves them
FigList = findobj(allchild(0), 'flat', 'Type', 'figure');
for iFig = 1:length(FigList)
    FigHandle = FigList(iFig);
    FigName   = get(FigHandle, 'Name');
    try
        savefig(FigHandle, fullfile(FolderName,['fig', num2str(iFig), '.fig']));
    catch
    end
end
```
