## Supplementary Zip file for "Mechanistic model for human brain metabolism and its connection to the neurovascular coupling": MetCost.m

```
function [f,c,gf,gc] = MetCost(theta,Met_Data,BOLD_data,modelName,amiOptions,ShouldIPlot,FID)

datasetIndex = [1,2,3,4];

negTheta = theta;
negTheta([22:23,2,31,34]) = theta([end-4:end]);
logL=0;
dlogL = zeros(numel(theta),1);
k = [10 10 10 10];

%% Steady state simulation
SteadyStateModel = str2func(['simulate_',modelName,'_SS']);
sol_SS = SteadyStateModel(inf,theta(1:57),k,[],amiOptions);

amiOptions.x0=sol_SS.x(end,:)';
k(5:9)=sol_SS.y(end,:);


DoubleStimModel = str2func(['simulate_',modelName,'_Double']);
SingleStimModel = str2func(['simulate_',modelName,'_Single']);
BOLDStimModel = str2func(['simulate_',modelName,'_BOLD']);

HuberStimulation = str2func(['simulate_',modelName,'_SNR']);

Lundengard2016_1secIPI=str2func(['simulate_',modelName,'_Lundengard2016_1secIPI']);
Lundengard2016_4secIPI=str2func(['simulate_',modelName,'_Lundengard2016_4secIPI']);

%% If gradient is to be calculated, 
if (nargout >2)
    %Settings for gradient
    amiOptions.sensi = 1;
    amiOptions.sensi_meth='forward';

    %Simulations
    sol_Double = DoubleStimModel(Met_Data(1).t,theta(1:57),k,Met_Data(1),amiOptions);
    sol_Single = SingleStimModel(Met_Data(2).t,theta(1:57),k,Met_Data(2),amiOptions);
    sol_BOLD = BOLDStimModel(BOLD_data(1).t,theta(1:57),k,BOLD_data(1),amiOptions);
    sol_negBOLD = BOLDStimModel(BOLD_data(1).t,negTheta(1:57),k,BOLD_data(2),amiOptions);
          
    simulationStatus = [sol_Double.status<0,sol_Single.status<0, sol_BOLD.status<0 ,sol_negBOLD.status<0];
    simulationChi2 = [sol_Double.chi2, sol_Single.chi2, sol_BOLD.chi2, sol_negBOLD.chi2];
    simulationSllh = [sol_Double.sllh, sol_Single.sllh, sol_BOLD.sllh, sol_negBOLD.sllh];
    %% if simulation is not sucessful
    if all(simulationStatus(datasetIndex))
        logL  = 1e98;
        dlogL = NaN(numel(theta),1);
    else
        %Calculate cost (done here automatically in simulation as
        %experimental data D is sent in to the simulation

        logL  = logL + sum(simulationChi2(datasetIndex));
        %Gradient
        dlogL = dlogL - sum(simulationSllh(datasetIndex));
    end
else % IF no gradient is wanted by optimization alg
    amiOptions.sensi=0;
    
    %Simulations
    sol_Double = DoubleStimModel(Met_Data(1).t,theta(1:57),k,Met_Data(1),amiOptions);
    sol_Single = SingleStimModel(Met_Data(2).t,theta(1:57),k,Met_Data(2),amiOptions);
    sol_BOLD = BOLDStimModel(BOLD_data(1).t,theta(1:57),k,BOLD_data(1),amiOptions);
    sol_negBOLD = BOLDStimModel(BOLD_data(1).t,negTheta(1:57),k,BOLD_data(2),amiOptions);
%     sol_HuberBOLD = HuberStimulation(HuberData.t,[theta(1:57);log10(30)],k,HuberData,amiOptions);
    
%     sol_Lundengard2016_1secIPI = Lundengard2016_1secIPI([],theta(1:57),k,lundengardData(1),amiOptions);
%     sol_Lundengard2016_4secIPI = Lundengard2016_4secIPI([],theta(1:57),k,lundengardData(2),amiOptions);
    
    simulationStatus = [sol_Double.status<0,sol_Single.status<0, sol_BOLD.status<0 ,sol_negBOLD.status<0];
    simulationChi2 = [sol_Double.chi2, sol_Single.chi2, sol_BOLD.chi2, sol_negBOLD.chi2];
    % see above
    if all(simulationStatus(datasetIndex))
        logL = 1e98;
    else 
        logL  = logL + sum(simulationChi2(datasetIndex));
    end
end

%% Return arguments
f=logL;
c=[]; % optimization-specific to eSS, if there are any constraints
gf=dlogL;
gc=[];% optmization-specific to eSS, gradient of potential constraints.

%% MCMC related stuff
   if nargin == 9 && logL < chi2inv(0.95,(2*28+8*24)) % Save to file. Here we had 248 datapoints, change to whatever data your're using.
        fprintf(FID,'%4.10f %10.10f ',[f, theta']); fprintf(FID,'\n');
   end
%% Plotting
if nargin>6 && ShouldIPlot==1
%     sol_HuberBOLD = HuberStimulation([HuberData.t(1):0.001:HuberData.t(end)],[theta(1:57),log10(30)],k,[],amiOptions);
    
    text={'Lactate','Glutamate','Glucose','Aspartate'};
    lims = {[-30,50],[-2,6],[-70,20],[-25,10];[-40,60],[-3,7],[-60,20],[-25,10]};
    stimTimes = {[594 1188] [1782 2376];[396 1188],[nan nan]};
    titles = {'Double Stimulus', 'Single stimulus'};
    sols = {sol_Double,sol_Single};
    
    for j = 1:2
        figure();
        for i = 1:4
            ax(i,j) = subplot(2,2,i);
            hold on
            errorbar(ax(i,j),Met_Data(j).t,Met_Data(j).Y(:,i),Met_Data(j).Sigma_Y(:,i),'r*');
            plot(sols{j}.t,sols{j}.y(:,i),'b','LineWidth',2);
            ax(i,j).YLabel.String = 'Response (\Delta%)';
            ax(i,j).XLabel.String = 'Time [s]';
            ax(i,j).Title.String = text{i};
            set(ax(i,j),{'YLim'},lims(j,i));
            line(ax(i,j),stimTimes{j,1},repmat(lims{j,i}(1),1,2),'LineWidth',5,'Color',[0 0 0]);
            line(ax(i,j),stimTimes{j,2},repmat(lims{j,i}(1),1,2),'LineWidth',5,'Color',[0 0 0]);
            hold off
        end
        suplabel(titles{j},'t');
    end
    % positive BOLD
    figure();
    hold on
    errorbar(BOLD_data(1).t(2:end),BOLD_data(1).Y(2:end,5),BOLD_data(1).Sigma_Y(2:end,5),'r*');
    plot(sol_BOLD.t,sol_BOLD.y(:,5),'b','LineWidth',2);
    line([0 0.5],[-0.5 -0.5],'LineWidth',5,'Color',[0 0 0]);
    title('Positive BOLD-response')
    % Negative BOLD 
    hold off
    figure();
    hold on
    errorbar(BOLD_data(2).t(2:end),BOLD_data(2).Y(2:end,5),BOLD_data(2).Sigma_Y(2:end,5),'r*');
    plot(sol_negBOLD.t,sol_negBOLD.y(:,5),'b','LineWidth',2);
    line([0 0.5],[-0.5 -0.5],'LineWidth',5,'Color',[0 0 0]);
    title('Negative BOLD-response')
    hold off
    
  
end
end
```
