## Supplementary Zip file for "Mechanistic model for human brain metabolism and its connection to the neurovascular coupling": NVC_Metabolism_model_V6_Bednarik_syms.m

```
function [model] = NVC_Metabolism_model_V6_Bednarik_syms()

% set the parametrisation of the problem options are 'log', 'log10' and
% 'lin' (default).
%am_stepfun(t,1000.5,1,1001,0)*
%
model.param = 'log10';

%%
% STATES

% create state syms
syms glut GABA Ca AA constrict delay1con delay2con delay3con delay1dil delay2dil delay3dil dilate oHb dHb O2_c O2_t glucose_c glucose_t glucoseDelay pyr lac OG OAA Asp Glu Gln

% create state vector
model.sym.x = [glut, GABA, Ca, AA, constrict, delay1con, delay2con, delay3con delay1dil, delay2dil, delay3dil dilate, oHb, dHb, O2_c O2_t glucose_c glucose_t glucoseDelay pyr, lac, OG, OAA, Asp, Glu, Gln];

%% 
% PARAMETERS ( for these sensitivities will be computed )

% create parameter syms
syms ky kmetabolic k1f k1b k_basal kflowBasal proportion1 proportion2 kO2bbb kGlucf kGlucb kBV kmgluc kGD PL CaBas sinkGABA sinkGlu sinkA sinkCon sinkDil kglut kgaba k3 k4 k5 k7 b3 b4 kdelay1d kdelay2d kdelay3d kdelay1c kdelay2c kdelay3c  Vmax_pyr1 Km_pyr2 k1 ky1 ky2 Vmax_OAA1 Km_OAA2 Vmax_OG1 Vmax_OG2 Vmax_Glu1 Vmax_Glu2 Vmax_Gln Vmax_Asp Vmax_PO ky3 ky4 Vmax_pyr3 kstim kstim2 Km_pyr3 k_volumeScale k_volumeScale2

% create parameter vector 
model.sym.p = [ky, kmetabolic, k1f, k1b, k_basal, kflowBasal, proportion1, proportion2, kO2bbb, kGlucf, kGlucb, kBV, kmgluc, kGD, PL, CaBas, sinkGABA, sinkGlu, sinkA, sinkCon, sinkDil, kglut, kgaba, k3, k4, k5, k7, b3, b4, kdelay1d, kdelay2d, kdelay3d, kdelay1c, kdelay2c, kdelay3c, Vmax_pyr1, Km_pyr2, k1, ky1, ky2, Vmax_OAA1, Km_OAA2, Vmax_OG1, Vmax_OG2, Vmax_Glu1, Vmax_Glu2, Vmax_Gln, Vmax_Asp, Vmax_PO, ky3, ky4, Vmax_pyr3, kstim, kstim2, Km_pyr3,k_volumeScale, k_volumeScale2];

% set the parametrisation of the problem options are 'log', 'log10' and
% 'lin' (default).

%% NVC CONSTANTS ( for these no sensitivities will be computed )
% this part is optional and can be ommited

% create parameter syms
syms Gbody O2body oHbbody dHbbody ssLac ssGlu ssGlucose_t ssAsp ssBOLD

% create parameter vector 
model.sym.k = [Gbody, O2body, oHbbody,dHbbody ssLac ssGlu ssGlucose_t ssAsp ssBOLD];

%% NVC SYSTEM EQUATIONS

% create symbolic variable for time
syms t
model.sym.xdot = sym(zeros(size(model.sym.x)));
syms CBV FlowVar
FlowVar=(kflowBasal+exp(b3*dilate-b4*constrict));
CBV=FlowVar^kBV;

% stimulus = am_stepfun(t,0.001,1,0.5,0); % fMRI stimulus
% stimulus = am_stepfun(t,396,1,1188,0); % Single stimulus
% stimulus = am_stepfun(t,594,1,1188,0)+am_stepfun(t,1782,1,2376,0); % double stimulus
stimulus = am_stepfun(t,318,1,636,0)+am_stepfun(t,954,1,1272,0); %Bednarik stimulus

model.sym.xdot(1) = stimulus*kglut-glut*sinkGlu;
model.sym.xdot(2) = stimulus*kgaba-GABA*sinkGABA;
model.sym.xdot(3) = (CaBas*(1+k3*glut)/(1+(GABA/k4)))-Ca*sinkA;
model.sym.xdot(4) = PL*Ca-(k5+k7)*AA;
model.sym.xdot(5) = kdelay3c*delay3con-constrict*sinkCon;
model.sym.xdot(6) = k5*AA-delay1con*kdelay1c;
model.sym.xdot(7) = delay1con*kdelay1c-kdelay2c*delay2con;
model.sym.xdot(8) = delay2con*kdelay2c-kdelay3c*delay3con;
model.sym.xdot(9) = k7*AA-kdelay1d*delay1dil;
model.sym.xdot(10) = kdelay1d*delay1dil-kdelay2d*delay2dil;
model.sym.xdot(11) = kdelay2d*delay2dil-kdelay3d*delay3dil;
model.sym.xdot(12) = delay3dil*kdelay3d-dilate*sinkDil;
model.sym.xdot(13) = k1b*dHb*O2_c-k1f*oHb+oHbbody*FlowVar-oHb*FlowVar;
model.sym.xdot(14) = -k1b*dHb*O2_c+k1f*oHb+dHbbody*FlowVar-dHb*FlowVar;
model.sym.xdot(15) = -k1b*dHb*O2_c+k1f*oHb+O2body*FlowVar-O2_c*FlowVar-kO2bbb*(O2_c-O2_t);
model.sym.xdot(16) = CBV*(kO2bbb*(O2_c-O2_t))-((k_basal*glucose_t*O2_t^proportion1)*proportion1-(glucoseDelay*glucose_t*O2_t^proportion2)*proportion2)*k_volumeScale2;
model.sym.xdot(17) = Gbody*FlowVar-glucose_c*FlowVar-(kGlucf*glucose_c/(kmgluc+glucose_c))+kGlucb*(glucose_t-glucose_c);
model.sym.xdot(18) = ((((kGlucf*glucose_c/(kmgluc+glucose_c))-kGlucb*(glucose_t-glucose_c))*CBV))*k_volumeScale-(k_basal*glucose_t*O2_t^proportion1)-(glucoseDelay*glucose_t*O2_t^proportion2);
model.sym.xdot(19) =  stimulus*kmetabolic-glucoseDelay*kGD;

%% Metabolism SYSTEM EQUATIONS
% v0 = u
% v1 = ((Vmax_glcv*glcv))
% v2 = ((Vmax_glcn*glcn))

v3 = ((Vmax_pyr1*pyr));
v4 = k1*lac;
vTCA1 = Vmax_PO*((pyr/(Km_pyr2+pyr))*(OAA/(Km_OAA2+OAA)));
vTCA2 = ((Vmax_OG1*OG));
vx1 = ((Vmax_OG2*OG));
vx2 = ((Vmax_OAA1*OAA));
vGln = ((Vmax_Glu1*Glu));
vGlu = ((Vmax_Gln*Gln));
v5 = ((Vmax_Glu2*Glu));
v6 = ((Vmax_Asp*Asp));
vpc = Vmax_pyr3*(pyr/(Km_pyr3+pyr));


model.sym.xdot(20)=(k_basal*glucose_t*O2_t^proportion1) + (glucoseDelay*glucose_t*O2_t^proportion2) -vTCA1-v3-vpc;
model.sym.xdot(21)=((v3-v4));
model.sym.xdot(22)=((vTCA1-vTCA2-vx1));
model.sym.xdot(23)=((vTCA2-vTCA1-vx2+vpc));
model.sym.xdot(24)= ((vx2-v6));
model.sym.xdot(25)=((vx1+vGlu-vGln-v5));
model.sym.xdot(26)=((vGln-vGlu));

%% NVC INITIAL CONDITIONS

model.sym.x0 = sym(zeros(size(model.sym.x)));

model.sym.x0(3) = 10;
model.sym.x0(4) = 10;
model.sym.x0(13) = 200;
model.sym.x0(14) = 100;
model.sym.x0(15) = 100;
model.sym.x0(16) = 200;
model.sym.x0(17) = 100;
model.sym.x0(18) = 200;
% OBSERVALES

model.sym.y = sym(zeros(1,5));

model.sym.y(5) = 100*((exp(-ky*dHb)-ssBOLD)/ssBOLD);

%% Metabolism INITIAL CONDITIONS

% model.sym.x0(1)=10;
model.sym.x0(20:26)=1;
% OBSERVALES

model.sym.y(1)=100*((ky1*lac-ssLac)./ssLac);
model.sym.y(2)=100*((ky2*Glu-ssGlu)./ssGlu);
model.sym.y(3)=100*((ky3*glucose_t-ssGlucose_t)./ssGlucose_t);
model.sym.y(4)=100*((ky4*Asp-ssAsp)./ssAsp);

end
```
