## Supplementary Zip file for "Mechanistic model for human brain metabolism and its connection to the neurovascular coupling": PlotResults.m

```
load('./Parameters/ParametersAndsimulations.mat');
%% load Data
[metData,BOLD_data,bednarikData,gruetterData] = GenerateData(); % Get the metabolite and BOLD data

nTimePoints = 1000;
timeVectors = [linspace(metData(1).t(1),metData(1).t(end),nTimePoints).',linspace(metData(2).t(1),metData(2).t(end),nTimePoints).',linspace(BOLD_data(1).t(1),BOLD_data(1).t(end),nTimePoints).',linspace(bednarikData.t(1),bednarikData.t(end),nTimePoints).',linspace(gruetterData.t(1),gruetterData.t(end),nTimePoints).',linspace(0,1000,nTimePoints).'];

%% Plots 
lineWidth = 1; %set the line width of the errorbars in all figures

%% Lin et. al. fit 
 titleText={'Lactate','Glutamate','Glucose','Aspartate'};
    lims = {[-30,50],[-2,6],[-70,20],[-25,10];[-40,60],[-3,7],[-60,20],[-25,10]};
    stimTimes = {[594 1188] [1782 2376];[396 1188],[nan nan]};
    titles = {'Double Stimulus', 'Single stimulus'};
%     sols = {sol_Double,sol_Single};
    
    for j = 1:2
        f1(j)=figure('units','normalized','outerposition',[0 0 1 1]);
        for i = 1:4
            ax(i,j) = subplot(2,2,i);
            hold on
            eBarHandle = errorbar(ax(i,j),metData(j).t,metData(j).Y(:,i),metData(j).Sigma_Y(:,i),'r*','LineWidth',lineWidth);
            %             plot(sols{j}.t,sols{j}.y(:,i),'b','LineWidth',2);
            fillHandle = fill([timeVectors(:,j);flipud(timeVectors(:,j))],[simulation_lb(:,i+(j-1)*4);flipud(simulation_ub(:,i+(j-1)*4))],'b','FaceAlpha',0.2);
            ax(i,j).YLabel.String = 'Response (\Delta%)';
            ax(i,j).XLabel.String = 'Time [s]';
            ax(i,j).Title.String = titleText{i};
            set(ax(i,j),{'YLim'},lims(j,i));
            line(ax(i,j),stimTimes{j,1},repmat(lims{j,i}(1),1,2),'LineWidth',5,'Color',[0 0 0]);
            line(ax(i,j),stimTimes{j,2},repmat(lims{j,i}(1),1,2),'LineWidth',5,'Color',[0 0 0]);
            hold off
            set(ax(i,j),'FontSize',22);
        end
        maximizeAxes(f1(j),2,2); %maximise the size of all subplots  
        legend(ax(2,j),ax(2,j).Children(end-1:end),{'Model Simulation Uncertainty','Experimental Data'})
        sgtitle(titles{j},'FontSize',26);
        setSubFigureNotations(ax(:,j),30); %add the A-D notations to the subfigures
    end
    
    %% BOLD data
    % positive BOLD fit
    f2 = figure('units','normalized','outerposition',[0 0 1 1]);
    ax2(1)= subplot(2,1,1);
    hold on
    errorbar(BOLD_data(1).t(2:end),BOLD_data(1).Y(2:end,5),BOLD_data(1).Sigma_Y(2:end,5),'r*','LineWidth',lineWidth);
    %     plot(sol_BOLD.t,sol_BOLD.y(:,5),'b','LineWidth',2);
    fill([timeVectors(:,3);flipud(timeVectors(:,3))],[simulation_lb(:,9);flipud(simulation_ub(:,9))],'b','FaceAlpha',0.2);
    line([0 0.5],[-0.5 -0.5],'LineWidth',5,'Color',[0 0 0]);
    title('Positive BOLD-response')
    set(ax2(1).YLabel,'String','Response (\Delta%)');
    set(ax2(1).XLabel,'String','Time [s]');
    set(ax2(1),{'FontSize','YLim'},{22,[-0.5 2.5]});
    hold off 
 
    legend(ax2(1),ax2(1).Children(end-1:end),{'Model Simulation Uncertainty','Experimental Data'});
     
    % Negative BOLD fit   
    ax2(2)= subplot(2,1,2);
    hold on
    errorbar(BOLD_data(2).t(2:end),BOLD_data(2).Y(2:end,5),BOLD_data(2).Sigma_Y(2:end,5),'r*','LineWidth',lineWidth);
    %     plot(sol_negBOLD.t,sol_negBOLD.y(:,5),'b','LineWidth',2);
    fill([timeVectors(:,3);flipud(timeVectors(:,3))],[simulation_lb(:,10);flipud(simulation_ub(:,10))],'b','FaceAlpha',0.2);
    line([0 0.5],[-0.5 -0.5],'LineWidth',5,'Color',[0 0 0]);

    title('Negative BOLD-response')
    set(ax2(2).YLabel,'String','Response (\Delta%)');
    set(ax2(2).XLabel,'String','Time [s]');
    set(ax2(2),{'FontSize','YLim'},{22,[-0.5 0.2]});
    hold off
    maximizeAxes(f2,2,1); %maximise the size of all subplots 

    setSubFigureNotations(ax2,30); %add the A-B notations to the subfigures
    %% Bednarik data Predicitons
    
    lims ={[-20,40],[-3,6],[-50,40],[-20,10]}; 
    f4=figure('units','normalized','outerposition',[0 0 1 1]);
    for i = 1:4
        ax4(i) = subplot(2,2,i);
        hold on
        errorbar(bednarikData.t,bednarikData.Y(:,i),bednarikData.Sigma_Y(:,i),'r*','LineWidth',lineWidth);
        fill([timeVectors(:,4);flipud(timeVectors(:,4))],[simulation_lb(:,i+10);flipud(simulation_ub(:,i+10))],'b','FaceAlpha',0.2);
        
        set(gca,'YLim',lims{i})
        line([318 636],repmat(lims{i}(1),1,2),'LineWidth',5,'Color',[0 0 0]);
        line([954 1272],repmat(lims{i}(1),1,2),'LineWidth',5,'Color',[0 0 0]);
        hold off
        title(titleText{i})
        ax4(i).YLabel.String = 'Response (\Delta%)';
        ax4(i).XLabel.String = 'Time [s]';
        set(ax4,'FontSize',22);
    end
    %     sgtitle('Bednarik data')
    maximizeAxes(f4,2,2); %maximise the size of all subplots 
    setSubFigureNotations(ax4,30); %add the A-D notations to the subfigures
    legend(ax4(2),ax4(2).Children(end-1:end),{'Model Prediction Uncertainty','Experimental Data'})
    %% Gruetter Data Predictions
    lims = {[-40,80],[-6,6]};
    f5=figure('units','normalized','outerposition',[0 0 1 1]);
    for i = 1:2
        ax5(i) = subplot(2,1,i);
        hold on
        errorbar(gruetterData.t,gruetterData.Y(:,i),gruetterData.Sigma_Y(:,i),'r*','LineWidth',lineWidth);
        fill([timeVectors(:,5);flipud(timeVectors(:,5))],[simulation_lb(:,i+14);flipud(simulation_ub(:,i+14))],'b','FaceAlpha',0.2);
        
        set(gca,'YLim',lims{i})
        line([300 600],repmat(lims{i}(1),1,2),'LineWidth',5,'Color',[0 0 0]);
        line([900 1200],repmat(lims{i}(1),1,2),'LineWidth',5,'Color',[0 0 0]);
        hold off
        title(titleText{i})
        ax5(i).YLabel.String = 'Response (\Delta%)';
        ax5(i).XLabel.String = 'Time [s]';
        set(ax5,'FontSize',22);
    end
%     sgtitle('Gruetter data')
    maximizeAxes(f5,2,1); %maximise the size of all subplots
    setSubFigureNotations(ax5,30); %add the A-D notations to the subfigures
    legend(ax5(1),ax5(1).Children(end-1:end),{'Model Prediction Uncertainty','Experimental Data'})
    
    %% lundengård 2016 predictions
    titleText = [titleText,{'BOLD'}];
    noDataPredictionTitles = {'Lundengard 2016 1 sec IPI','Lundengard 2016 4 sec IPI'};
    load('.\Data\lundengardData.mat')
    
    for j = 1:2
        figure('units','normalized','outerposition',[0 0 1 1]);
        for i = 1:5
            ax6(i,j) = subplot(3,2,i);
            hold on
            fill([timeVectors(:,6);flipud(timeVectors(:,6))],[simulation_lb(:,i+(16+(j-1)*5));flipud(simulation_ub(:,i+(16+(j-1)*5)))].*8.7,'b','FaceAlpha',0.2);
            
            
            line([0 0.5],repmat(ax6(i,j).YLim(1),1,2),'LineWidth',5,'Color',[0 0 0]);
            line([1.5+(j-1)*3 5],repmat(ax6(i,j).YLim(1),1,2),'LineWidth',5,'Color',[0 0 0]);
            title(titleText{i})
            ax6(i,j).YLabel.String = 'Response (\Delta%)';
            ax6(i,j).XLabel.String = 'Time [s]';
            
            if i==5
                errorbar(lundengardData(j).t,lundengardData(j).Y(:,end),lundengardData(j).Sigma_Y(:,end),'r*')
            end
           set(ax6(i,j),'XLim',[0 100]);
        end
        set(ax6(5,j),'XLim',[0 30]);
%     sgtitle(noDataPredictionTitles{j})    
    end
    
    set(ax6,'FontSize',22);
    f=gcf;
%     maximizeAxes(f,3,2)
    setSubFigureNotations(ax6(:,2),30); %add the A-D notations to the subfigures
    %%
modelName = 'NVC_Metabolism_model_V6'; % set model Name

SNRModel = str2func(['simulate_',modelName,'_SNR']);

%set the time vectors for simulation such that all simualtions have the same number of elements
timeVector = [[0:0.1:1],logspace(0,4,nTimePoints)].';

%% simulation
tStimEnd = [[0.5:0.1:0.9],[1:10],[20:10:100],[120:20:580],[600:120:1200],5e3,1e4];

load('./Results/stimTime2SigalAmplitude.mat');
count = size(tStimEnd,2)+1;

simulation_UB = cellfun(@max,allSimulations,repmat({[]},1,count-1),num2cell(3*ones(1,count-1)),'UniformOutput',false); % get the maximum simulation value 
simulation_LB = cellfun(@min,allSimulations,repmat({[]},1,count-1),num2cell(3*ones(1,count-1)),'UniformOutput',false); % get the minimum simulation value 
%% find SNR

[rowIdx,colIdx] = find(min(signalAmplitude(:,[1:4],:),[],3)>2*mean(metData(1).Sigma_Y(:,[1:4])));
[~,tempIdx]=unique(colIdx);

%% plot max absolute amplitude vs stimulation time

f1=figure('units','normalized','outerposition',[0 0 1 1]);
% colour = linspecer(size(sol_SNR.y,2));

colour =[0.5  0.05 0.05
         0.95 0.53 0.13
         0    0.40 1.00
         0    0.40 0.20
         0.80 0    0
         0    0    1
         0.60 0    0.80
         1.00 0.12 0.36
         0.49 0.18 0.56
         0    0.60 0.87
         0    0.80 0.43
         0.69 0.35 0.73
         1.00 0.77 0.11
         1.00 0.70 0.58
         0.63 0.70 0.73
         0.64 0.47 0.11
         ];
     
AlphaLevel = 0.3; % set the alpha level for shaded areas in plots

ax1 = subplot(3,1,1);
hold on
timePoints2Plot_Idx = find(ismember(tStimEnd,[5 20 60 120 300 600 1200]));
xShift = [0.03 0.03 0.09 0.09 0.15 0.15 0.15 0.15];
% add lines at specific time points
for i = 1: size(timePoints2Plot_Idx,2)
   line(repmat(tStimEnd(timePoints2Plot_Idx(i)),2,1).',[0 55],'color',(colour(i,:)+(1-colour(i,:))*(1-AlphaLevel)),'LineWidth',5)% plota a line at each time pointe in the corresponding coulour
   text(tStimEnd(timePoints2Plot_Idx(i))-xShift(i),57.7,[num2str(tStimEnd(timePoints2Plot_Idx(i))),' s'],'FontWeight','bold','FontSize',32,'HorizontalAlignment','center'); %add the text lable at the time points
end

% plot max response curves
for i = 1:size(sol_SNR{1}(1).y,2)-2
    fill([tStimEnd.';flipud(tStimEnd.')],[min(signalAmplitude(:,i,:),[],3);flipud(max(signalAmplitude(:,i,:),[],3))],'b','FaceColor',(colour(end-1,:)+(1-colour(end-1,:))*(1-0.5)),'FaceAlpha',1)
    text(2e3,round(max(max(signalAmplitude(:,i,:),[],3)))+2,titleText{i},'FontSize',36,'FontWeight','bold'); %add the text label for each metabolite
end
% plot(tStimEnd,signalAmplitude(:,[1:4],1),'*','MarkerSize',10,'LineWidth',2);

% hleg = legend(titleText,'FontSize',32);
% hleg.AutoUpdate='Off'; %prevet Legend to auto add new lines
xlabel('Length of Stimulation [s]','FontSize',32)
ylabel(sprintf('Maximal signal amplitude\n relative baseline'),'FontSize',32)
set(ax1,{'XScale','XLim','FontSize','YTickLabels'},{'log',[0.5 1e4],32,strcat(ax1.YTickLabels,' %')});

maximizeAxes(f1,1,1);
setSubFigureNotations(ax1,32); %add the A-B notations to the subfigures
%% plot responses
f2=figure('units','normalized','outerposition',[0 0 1 1]);


count = 1;
for j = timePoints2Plot_Idx
    
    for i = 1:4
        ax(i) = subplot(3,2,i+2);
        hold on
        fill([timeVector;flipud(timeVector)],[simulation_LB{j}(:,i);flipud(simulation_UB{j}(:,i))],'b','FaceColor',colour(count,:),'FaceAlpha',AlphaLevel);

        title(titleText{i})
        ax(i).YLabel.String = 'Response (\Delta%)';
        ax(i).XLabel.String = 'Time [s]';
        set(ax(i),{'FontSize','XLim'},{32,[0 2000]});
    end

    count = count+1;
end
legend(ax(2),cellfun(@sprintf,repmat({'%d s'},1,size(timePoints2Plot_Idx,2)),num2cell(tStimEnd(timePoints2Plot_Idx)),'Uni',0),'Orientation','horizontal')
maximizeAxes(f2,2,2);
setSubFigureNotations(ax,32); %add the A-B notations to the subfigures
%% Parameter histograms
parameterNames = {'ky', 'kmetabolic', 'k1f', 'k1b', 'k_basal', 'kflowBasal', 'proportion1', 'proportion2', 'kO2bbb', 'kGlucf', 'kGlucb', 'kBV', 'kmgluc', 'kGD', 'PL', 'CaBas', 'sinkGABA', 'sinkGlu', 'sinkA', 'sinkCon', 'sinkDil', 'kglut', 'kgaba', 'k3', 'k4', 'k5', 'k7', 'b3', 'b4', 'kdelay1d', 'kdelay2d', 'kdelay3d', 'kdelay1c', 'kdelay2c', 'kdelay3c', 'Vmax_pyr1', 'Km_pyr2', 'k1', 'ky1', 'ky2', 'Vmax_OAA1', 'Km_OAA2', 'Vmax_OG1', 'Vmax_OG2', 'Vmax_Glu1', 'Vmax_Glu2', 'Vmax_Gln', 'Vmax_Asp', 'Vmax_PO', 'ky3', 'ky4', 'Vmax_pyr3', 'kstim', 'kstim2', 'Km_pyr3','k_volumeScale','k_volumeScale2'};
parameterNames = [parameterNames, strcat('Neg_',parameterNames([22:23,2,31,34]))];

%Load parameters
load('./Parameters/opt(213.44)_201001-034806_v6.mat')

figure()
for i=1:size(allParameters,1)
subplot(8,8,i)
h=histogram(10.^(allParameters(i,:)));
hold on
line(repmat(10.^(optParamM(i,1)),1,2),[0 max(h.Values)],'Color','r','LineWidth',2);
title(parameterNames{i})
end
```
