## Supplementary Zip file for "Mechanistic model for human brain metabolism and its connection to the neurovascular coupling": Run.m

```
clear all
close all
clc

warning 'off'

addpath('..');


msg = "";
opts = ["Plot Figures" ....
        "Run simulations"
        "Run parameter sampling"
        "Run parameter estimation"
        "Generate models"
        ];

choice = menu(msg,opts);
addpath(genpath(fullfile(pwd,'Matlab Files')))
try

    switch choice
        case 1
           PlotResults
        case 2
            simulateAndPlotMCMCresult
        case 3
            MCMC_sampling
        case 4
            runParallelOpt
        case 5
            version = ver('MATLAB');
            year = str2double(version.Release(3:6));
            if year > 2017
                ver('MATLAB')
                throw(MException('ME:MatlabVersion', 'The Generate Models function uses AMICI''s amiwrap function and does not work with Matlab versions newer than Matlab r2017b \n'));
            end
            GenerateModels
    end
catch err
   close all 
   disp(err)
   disp("The systems biology toolbox (http://www.sbtoolbox2.org/main.php) is necessary to run the scripts in this project.\n")
   disp("It must be compiled and added to the MATLAB path before the scripts are run.")
end
```
