## Supplementary Zip file for "Mechanistic model for human brain metabolism and its connection to the neurovascular coupling": runParallel.sh

```
#!/bin/bash
#
# Use 2 nodes, i.e. 2x64 cores, for 1 hour
#SBATCH -N 2
#SBATCH -t 09:30:00
#SBATCH -A LiU-2019-16
#SBATCH --exclusive

# Update the project if possible. 
#git pull

modelName='NVC_Metabolism_model_V6'
cdprojpath="cd('/proj/13cmfabrain/x_nicsu/brain_Projects/NVC_Metabolism_Model_AMICI_V2');"

#Use hidden matlab r2017 module to compile AMICII models
module load MATLAB/.R2017b-nsc1
MATLAB_0='matlab -nodesktop -nodisplay -singleCompThread'
${MATLAB_0} -r "GenerateModels('${modelName}'); exit"

# Load the Matlab module
module load MATLAB/R2020a-nsc1 
module load parallel/20181122-nsc1
#module add gcc/8.2.0 

MATLAB='matlab -nodesktop -nodisplay -singleCompThread'
# The name of the Matlab script (without .m)
#job=runParameterEstimation_v2
job=runParameterEstimation_v2

#save_valid=0
now=$(date '+%Y%m%d-%H%M%S')

# Setup things first. 
${MATLAB} "\"$cdprojectpath Setup('${now}'); exit\""

# How many srun processes to keep running. 
MAX_SRUN=$(hostlist -e --repeat-slurm-tasks=$SLURM_TASKS_PER_NODE $SLURM_JOB_NODELIST | wc -l)


# Run main task. Note the explicit "exit" to exit Matlab.
# "Using GNU Parallel to launch multiple tasks but no more than $MAX_SRUN at any one time"
#seq 1 1500| parallel -j ${MAX_SRUN} "srun -N1 -n1  ${MATLAB} -r "'"runParameterEstimation_v2($RANDOM,'"'${now}'"','"'${modelName}'"','${save_valid}',0);exit"'" > ./Log/${job}_{}.log" 

# Run main task. Note the explicit "exit" to exit Matlab.
seq 1 496| parallel --ssh=jobsh -S $(hostlist -e -s',' -d  -p "$SLURM_CPUS_ON_NODE/" $SLURM_JOB_NODELIST) ${MATLAB} -r "\"$cdprojpath  ${job}($RANDOM,'${now}','${modelName}',{}); exit\""

# If you want the output from each Matlab instance in a separate file, you can e.g use "--results=output-${modelName}"
# If you want to watch the output as it is printed, you can use "--line-buffer".
# To see more of the internal workings of Parallel, add "-vv"
# Recommended reading: "man parallel"
# End of script
```
