## Supplementary Zip file for "Mechanistic model for human brain metabolism and its connection to the neurovascular coupling": runParallelOpt.m

```
% !git pull
folderName = datestr(now,'yyyymmdd-HHMMSS');

modelName = 'NVC_Metabolism_model_V6';
GenerateModels(modelName)


Setup(folderName)
rng('shuffle')

parfor i=1:1 %1:n, n = number of repeats
    random=randi([0,32767],1);
    runParameterEstimation_v2(random,folderName,modelName,0,1)
end
```
