## Supplementary Zip file for "Mechanistic model for human brain metabolism and its connection to the neurovascular coupling": runParameterEstimation_v2.m

```
function runParameterEstimation_v2(worker,folderName,modelName, save_valid, show_iter)
s = rng('shuffle');
rng(s.Seed+worker)

if nargin<2 || isempty(folderName)
    folderName='Temp';
end
%% set Paths
run('../Toolboxes/AMICI-0.10.11_SS_eventFix/matlab/installAMICI.m')% set path for the toolbox.
addpath(genpath(fullfile(pwd,'../Toolboxes/jrbanga_-meigo64-33e109c94b25/MEIGO'))) %add the MEIGO folder to the path

% set path for the toolbox.

addpath(genpath(fullfile(pwd,'Models')))
addpath(genpath(fullfile(pwd)))
%% load data

[metData,BOLD_data,~,~] = GenerateData(); % Get the metabolite and BOLD data
load('.\Data\lundengardData.mat')

%% set amiOptions
options=amioption();
options.maxsteps=1e4;
options.atol=1e-16;
options.rtol=1e-8;

%%

folder = sprintf('./Results/%s/%s/', folderName, modelName);
if ~exist(folder,'dir')
    mkdir(folder)
end

if nargin>2 && save_valid
    fid=fopen(sprintf('%s/validParams-%s-%i.csv', folder,datestr(now,'yymmdd-HHMMSS'),worker),'wt');
    fprintf(fid,['Cost,' strjoin(paramNames,', ') '\n']);
else
    fid=[];
end

%% MEIGO stuff
if isempty(dir(sprintf('./Results/%s/%s/*.mat', folderName, modelName)))
    %if the results folder is empty load previous parameters as start guess
    load('./Parameters/opt(213.44)_201001-034806_v6.mat')
    
else
    saveFolder = sprintf('./Results/%s/%s',folderName, modelName);
    bestParamFileName = findBestSavedParameters(saveFolder);
    load(sprintf('./Results/%s/%s/%s', folderName, modelName,bestParamFileName))
end
startGuess = optParamM; % startGuess

problem.x_L       = log10(1e-4*ones(size(startGuess)));
problem.x_U       = log10(7e4*ones(size(startGuess)));

% MEIGO OPTIONS I (COMMON TO ALL SOLVERS):
opts.ndiverse     = 500; %100; %500; %5; %
opts.maxtime      = 3600; % In cess this option will be overwritten
opts.maxeval      = 1e8;
opts.log_var      = [];%;[1:size(startGuess,2)]; 

opts.local.solver = 'dhc'; %'fmincon'; %'nl2sol'; %'mix'; %
opts.local.finish = opts.local.solver; 
opts.local.bestx = 0;
opts.local.n2=10;
opts.local.balance=0.2;
% problem.f         = 'Gluc_costFunction';
problem.f = 'MetCost';
problem.x_0 = startGuess;

% MEIGO OPTIONS II (FOR ESS AND MULTISTART):
opts.local.iterprint = 1;

% MEIGO OPTIONS III (FOR ESS ONLY):
opts.dim_refset   = 'auto'; %
optim_algorithm = 'ess'; % 'multistart'; %  'cess'; %

% OPTIONS AUTOMATICALLY SET AS A RESULT OF PREVIOUS OPTIONS:
if(strcmp(opts.local.solver,'fmincon'))
	opts.local.use_gradient_for_finish = 1; %DW: provide gradient to fmincon
else
	opts.local.use_gradient_for_finish = 0; %DW: provide gradient to fmincon 
end
opts.local.check_gradient_for_finish = 0; %DW: gradient checker

%%
warning('off','all')

Results = MEIGO(problem,opts,optim_algorithm,metData,BOLD_data,modelName,options,0); 
fitting_speed     = Results.time(end);
minfvalM           = Results.fbest;
optParamM    = Results.xbest'; 


save(sprintf('./Results/%s/%s/opt(%.2f)_%s.mat',folderName, modelName, minfvalM,  datestr(now,'yymmdd-HHMMSS'))  ,'optParamM') % Saves the best parameters found.

warning('on','all')
end
```
