## Supplementary Zip file for "Mechanistic model for human brain metabolism and its connection to the neurovascular coupling": Setup.m

```
function []=Setup(folderName)
%% Setup path and toolboxes
if ~exist('amiwrap','file')
% run('../Toolboxes/AMICI-master/matlab/installAMICI.m')
run('../Toolboxes/AMICI-0.10.11_SS_eventFix/matlab/installAMICI.m')
end

addpath(genpath(fullfile(pwd,'Models')))
addpath(genpath(fullfile(pwd)))
%% Create necessary folders and compile models
if ~exist('Log','dir')
    mkdir('Log')
end

if nargin<1 || isempty(folderName)
    location='./Results/Temp/';
else
    location=['./Results/' folderName '/'];
end


% GenerateModels

if ~exist(location,'dir')
mkdir(location)
end

end
```
