## Supplementary Zip file for "Mechanistic model for human brain metabolism and its connection to the neurovascular coupling": simulateAndPlotMCMCresult.m

```
%% set Paths
clear all 
close all


addpath(genpath(fullfile(pwd,'Models')))
addpath(genpath(fullfile(pwd,'Data')))
addpath(genpath(fullfile(pwd,'lib')))
addpath(genpath(fullfile(pwd)))

%% load Data
[metData,BOLD_data,bednarikData,gruetterData] = GenerateData(); % Get the metabolite and BOLD data
%% Here i've in a previous article specified sim options outside of costfunction, do as you wish
% set amiOptions
options=amioption();
options.maxsteps=1e4;
options.atol=1e-16;
options.rtol=1e-8;
amiOptions.sensi=0;

modelName = 'NVC_Metabolism_model_V6'; % set model Name

%Load parameters
load('./Parameters/opt(213.44)_201001-034806_v6.mat')

if isempty(dir('./Results/MCMCResult_*'))
fprintf('No MCMC sapling results found in Results folder! Run parameter sampling option first.')
return
end
MCMCResultFolders = dir('./Results/MCMCResult_*');
MCMCResultFile =  dir(sprintf('./Results/%s/MCMC_%s_*',MCMCResultFolders(1).name,modelName));
MCMCResults = load(sprintf('./Results/%s/%s',MCMCResultFolders(1).name,MCMCResultFile(1).name));

threshold = chi2inv(0.95,248-58);
allParameters = [optParamM,MCMCResults(MCMCResults(:,1)<threshold,2:end).'];


%setup model simulation
DoubleStimModel = str2func(['simulate_',modelName,'_Double']);
SingleStimModel = str2func(['simulate_',modelName,'_Single']);
BOLDStimModel = str2func(['simulate_',modelName,'_BOLD']);
BednarikModel = str2func(['simulate_',modelName,'_Bednarik']); 
GruetterModel =  str2func(['simulate_',modelName,'_Gruetter']);

Lundengard2016_1secIPI=str2func(['simulate_',modelName,'_Lundengard2016_1secIPI']);
Lundengard2016_4secIPI=str2func(['simulate_',modelName,'_Lundengard2016_4secIPI']);

nTimePoints = 1000;
%set the time vectors for simulation such that all simualtions have the same number of elements
timeVectors = [linspace(metData(1).t(1),metData(1).t(end),nTimePoints).',linspace(metData(2).t(1),metData(2).t(end),nTimePoints).',linspace(BOLD_data(1).t(1),BOLD_data(1).t(end),nTimePoints).',linspace(bednarikData.t(1),bednarikData.t(end),nTimePoints).',linspace(gruetterData.t(1),gruetterData.t(end),nTimePoints).',linspace(0,1000,nTimePoints).'];
simulations = nan(nTimePoints,size(allParameters,2),28);

%% simulate all paarameters
for i =1:size(allParameters,2)
    theta = allParameters(:,i);
    negTheta = theta;
    negTheta([22:23,2,31,34]) = theta([end-4:end]);
    
    k = [10 10 10 10];
    %% Steady state simulation
    SteadyStateModel = str2func(['simulate_',modelName,'_SS']);
    sol_SS = SteadyStateModel(inf,theta(1:57),k,[],amiOptions);
    amiOptions.x0=sol_SS.x(end,:)';
    k(5:9)=sol_SS.y(end,:);

    % Simulations
    sol_Double = DoubleStimModel(timeVectors(:,1),theta(1:57),k,[],amiOptions);
    sol_Single = SingleStimModel(timeVectors(:,2),theta(1:57),k,[],amiOptions);
    sol_BOLD = BOLDStimModel(timeVectors(:,3),theta(1:57),k,[],amiOptions);
    sol_negBOLD = BOLDStimModel(timeVectors(:,3),negTheta(1:57),k,[],amiOptions);
    
    %Data validation predictions
    sol_Bednarik = BednarikModel(timeVectors(:,4),theta(1:57),k,[],amiOptions);
    sol_Gruetter = GruetterModel(timeVectors(:,5),theta(1:57),k,[],amiOptions);
    
    % no-data Predictions
    sol_Lundengard2016_1secIPI = Lundengard2016_1secIPI(timeVectors(:,6),theta(1:57),k,[],amiOptions);
    sol_Lundengard2016_4secIPI = Lundengard2016_4secIPI(timeVectors(:,6),theta(1:57),k,[],amiOptions);
    
    
    
    simulations(:,i,:) = permute([sol_Double.y(:,1:4),sol_Single.y(:,1:4),sol_BOLD.y(:,5),sol_negBOLD.y(:,5),sol_Bednarik.y(:,1:4),sol_Gruetter.y(:,1:2),sol_Lundengard2016_1secIPI.y,sol_Lundengard2016_4secIPI.y,sol_Double.y(:,5),sol_Single.y(:,5)],[1,3,2]);
end
simulation_ub = permute(max(simulations,[],2),[1,3,2]);% get the max value for each simulated timepoint
simulation_lb = permute(min(simulations,[],2),[1,3,2]);%% get the min value for each simulated timepoint

%% Plots 
lineWidth = 1; %set the line width of the errorbars in all figures

%% Lin et. al. fit 
 text={'Lactate','Glutamate','Glucose','Aspartate'};
    lims = {[-30,50],[-2,6],[-70,20],[-25,10];[-40,60],[-3,7],[-60,20],[-25,10]};
    stimTimes = {[594 1188] [1782 2376];[396 1188],[nan nan]};
    titles = {'Double Stimulus', 'Single stimulus'};
%     sols = {sol_Double,sol_Single};
    
    for j = 1:2
        f1(j)=figure('units','normalized','outerposition',[0 0 1 1]);
        for i = 1:4
            ax(i,j) = subplot(2,2,i);
            hold on
            eBarHandle = errorbar(ax(i,j),metData(j).t,metData(j).Y(:,i),metData(j).Sigma_Y(:,i),'r*','LineWidth',lineWidth);
            %             plot(sols{j}.t,sols{j}.y(:,i),'b','LineWidth',2);
            fillHandle = fill([timeVectors(:,j);flipud(timeVectors(:,j))],[simulation_lb(:,i+(j-1)*4);flipud(simulation_ub(:,i+(j-1)*4))],'b','FaceAlpha',0.2);
            ax(i,j).YLabel.String = 'Response (\Delta%)';
            ax(i,j).XLabel.String = 'Time [s]';
            ax(i,j).Title.String = text{i};
            set(ax(i,j),{'YLim'},lims(j,i));
            line(ax(i,j),stimTimes{j,1},repmat(lims{j,i}(1),1,2),'LineWidth',5,'Color',[0 0 0]);
            line(ax(i,j),stimTimes{j,2},repmat(lims{j,i}(1),1,2),'LineWidth',5,'Color',[0 0 0]);
            hold off
            set(ax(i,j),'FontSize',22);
        end
        maximizeAxes(f1(j),2,2); %maximise the size of all subplots  
        legend(ax(2,j),ax(2,j).Children(end-1:end),{'Model Simulation Uncertainty','Experimental Data'})
        sgtitle(titles{j},'FontSize',26);
        setSubFigureNotations(ax(:,j),30); %add the A-D notations to the subfigures
    end
    
    %% BOLD data
    % positive BOLD fit
    f2 = figure('units','normalized','outerposition',[0 0 1 1]);
    ax2(1)= subplot(2,1,1);
    hold on
    errorbar(BOLD_data(1).t(2:end),BOLD_data(1).Y(2:end,5),BOLD_data(1).Sigma_Y(2:end,5),'r*','LineWidth',lineWidth);
    %     plot(sol_BOLD.t,sol_BOLD.y(:,5),'b','LineWidth',2);
    fill([timeVectors(:,3);flipud(timeVectors(:,3))],[simulation_lb(:,9);flipud(simulation_ub(:,9))],'b','FaceAlpha',0.2);
    line([0 0.5],[-0.5 -0.5],'LineWidth',5,'Color',[0 0 0]);
    title('Positive BOLD-response')
    set(ax2(1).YLabel,'String','Response (\Delta%)');
    set(ax2(1).XLabel,'String','Time [s]');
    set(ax2(1),{'FontSize','YLim'},{22,[-0.5 2.5]});
    hold off 
 
    legend(ax2(1),ax2(1).Children(end-1:end),{'Model Simulation Uncertainty','Experimental Data'});
     
    % Negative BOLD fit   
    ax2(2)= subplot(2,1,2);
    hold on
    errorbar(BOLD_data(2).t(2:end),BOLD_data(2).Y(2:end,5),BOLD_data(2).Sigma_Y(2:end,5),'r*','LineWidth',lineWidth);
    %     plot(sol_negBOLD.t,sol_negBOLD.y(:,5),'b','LineWidth',2);
    fill([timeVectors(:,3);flipud(timeVectors(:,3))],[simulation_lb(:,10);flipud(simulation_ub(:,10))],'b','FaceAlpha',0.2);
    line([0 0.5],[-0.5 -0.5],'LineWidth',5,'Color',[0 0 0]);

    title('Negative BOLD-response')
    set(ax2(2).YLabel,'String','Response (\Delta%)');
    set(ax2(2).XLabel,'String','Time [s]');
    set(ax2(2),{'FontSize','YLim'},{22,[-0.5 0.2]});
    hold off
    maximizeAxes(f2,2,1); %maximise the size of all subplots 

    setSubFigureNotations(ax2,30); %add the A-B notations to the subfigures
    %% Bednarik data Predicitons
    
    lims ={[-20,40],[-3,6],[-50,40],[-20,10]}; 
    f4=figure('units','normalized','outerposition',[0 0 1 1]);
    for i = 1:4
        ax4(i) = subplot(2,2,i);
        hold on
        errorbar(bednarikData.t,bednarikData.Y(:,i),bednarikData.Sigma_Y(:,i),'r*','LineWidth',lineWidth);
        fill([timeVectors(:,4);flipud(timeVectors(:,4))],[simulation_lb(:,i+10);flipud(simulation_ub(:,i+10))],'b','FaceAlpha',0.2);
        
        set(gca,'YLim',lims{i})
        line([318 636],repmat(lims{i}(1),1,2),'LineWidth',5,'Color',[0 0 0]);
        line([954 1272],repmat(lims{i}(1),1,2),'LineWidth',5,'Color',[0 0 0]);
        hold off
        title(text{i})
        ax4(i).YLabel.String = 'Response (\Delta%)';
        ax4(i).XLabel.String = 'Time [s]';
        set(ax4,'FontSize',22);
    end
    %     sgtitle('Bednarik data')
    maximizeAxes(f4,2,2); %maximise the size of all subplots 
    setSubFigureNotations(ax4,30); %add the A-D notations to the subfigures
    legend(ax4(2),ax4(2).Children(end-1:end),{'Model Prediction Uncertainty','Experimental Data'})
    %% Gruetter Data Predictions
    lims = {[-40,80],[-6,6]};
    f5=figure('units','normalized','outerposition',[0 0 1 1]);
    for i = 1:2
        ax5(i) = subplot(2,1,i);
        hold on
        errorbar(gruetterData.t,gruetterData.Y(:,i),gruetterData.Sigma_Y(:,i),'r*','LineWidth',lineWidth);
        fill([timeVectors(:,5);flipud(timeVectors(:,5))],[simulation_lb(:,i+14);flipud(simulation_ub(:,i+14))],'b','FaceAlpha',0.2);
        
        set(gca,'YLim',lims{i})
        line([300 600],repmat(lims{i}(1),1,2),'LineWidth',5,'Color',[0 0 0]);
        line([900 1200],repmat(lims{i}(1),1,2),'LineWidth',5,'Color',[0 0 0]);
        hold off
        title(text{i})
        ax5(i).YLabel.String = 'Response (\Delta%)';
        ax5(i).XLabel.String = 'Time [s]';
        set(ax5,'FontSize',22);
    end
%     sgtitle('Gruetter data')
    maximizeAxes(f5,2,1); %maximise the size of all subplots
    setSubFigureNotations(ax5,30); %add the A-D notations to the subfigures
    legend(ax5(1),ax5(1).Children(end-1:end),{'Model Prediction Uncertainty','Experimental Data'})
    
    %% lundengård 2016 predictions
    text = [text,{'BOLD'}];
    noDataPredictionTitles = {'Lundengard 2016 1 sec IPI','Lundengard 2016 4 sec IPI'};
    load('.\Data\lundengardData.mat')
    
    for j = 1:2
        figure('units','normalized','outerposition',[0 0 1 1]);
        for i = 1:5
            ax6(i,j) = subplot(3,2,i);
            hold on
            fill([timeVectors(:,6);flipud(timeVectors(:,6))],[simulation_lb(:,i+(16+(j-1)*5));flipud(simulation_ub(:,i+(16+(j-1)*5)))].*8.7,'b','FaceAlpha',0.2);
            
            
            line([0 0.5],repmat(ax6(i,j).YLim(1),1,2),'LineWidth',5,'Color',[0 0 0]);
            line([1.5+(j-1)*3 5],repmat(ax6(i,j).YLim(1),1,2),'LineWidth',5,'Color',[0 0 0]);
            title(text{i})
            ax6(i,j).YLabel.String = 'Response (\Delta%)';
            ax6(i,j).XLabel.String = 'Time [s]';
            
            if i==5
                errorbar(lundengardData(j).t,lundengardData(j).Y(:,end),lundengardData(j).Sigma_Y(:,end),'r*')
            end
           set(ax6(i,j),'XLim',[0 100]);
        end
        set(ax6(5,j),'XLim',[0 30]);
%     sgtitle(noDataPredictionTitles{j})    
    end
    
    set(ax6,'FontSize',22);
    f=gcf;

    setSubFigureNotations(ax6(:,2),30); %add the A-D notations to the subfigures
  

    
```
