## Supplementary Zip file for "Mechanistic model for human brain metabolism and its connection to the neurovascular coupling": stimTime2SigalAmplitude_analysis.m

```
%% set Paths
% clear all 
% close all
% 
% 
% run('../Toolboxes/AMICI-master/matlab/installAMICI.m')% set path for the toolbox.
% addpath(genpath(fullfile(pwd,'Models')))
% addpath(genpath(fullfile(pwd,'Data')))
% addpath(genpath(fullfile(pwd)))
% addpath(genpath(fullfile('../Toolboxes')))
%% load Data
% [metData,BOLD_data,bednarikData,gruetterData] = GenerateData(); % Get the metabolite and BOLD data
% %% Here i've in a previous article specified sim options outside of costfunction, do as you wish
% % set amiOptions
% options=amioption();
% options.maxsteps=1e4;
% options.atol=1e-16;
% options.rtol=1e-8;
% amiOptions.sensi=0;

modelName = 'NVC_Metabolism_model_V6'; % set model Name

%Load parameters
% load('./Parameters/opt(213.44)_201001-034806_v6.mat')
% MCMCResults=load('./Results/MCMCResult_210910_050734/MCMC_NVC_Metabolism_model_V6_210908_205112.dat');
% allParameters = [optParamM,MCMCResults];


SNRModel = str2func(['simulate_',modelName,'_SNR']);


nTimePoints = 1000;
%set the time vectors for simulation such that all simualtions have the same number of elements
timeVector = [[0:0.1:1],logspace(0,4,nTimePoints)].';

%% simulation
tStimEnd = [[0.5:0.1:0.9],[1:10],[20:10:100],[120:20:580],[600:120:1200],5e3,1e4];

if isempty(dir('./Results/stimTime2SigalAmplitude.mat')) %if no saved file exists 
    signalAmplitude = zeros(size(tStimEnd,2),6,size(allParameters,2));%cell(size(acceptableParameters,2),1);
    allSimulations = cell(1,size(tStimEnd,2));
    sol_SNR = cell(1,size(tStimEnd,2));
    count = 1;
    for i = tStimEnd
        for j = 1:size(allParameters,2)
            theta = [allParameters(:,j);log10(i)];
            negTheta = theta;
            negTheta([22:23,2,31,34]) = theta([end-5:end-1]);
            
            k = [10 10 10 10];
            %% Steady state simulation
            SteadyStateModel = str2func(['simulate_',modelName,'_SS']);
            sol_SS = SteadyStateModel(inf,theta(1:57),k,[],amiOptions);
            amiOptions.x0=sol_SS.x(end,:)';
            k(5:9)=sol_SS.y(end,:);
            
            % Simulations
            sol_SNR{count}(j) = SNRModel(timeVector,theta([1:57,end]),k,[],amiOptions);
            % plotStates(sol_SNR,theta)
            
            signalAmplitude(count,:,j) =max(abs(sol_SNR{count}(j).y))-sol_SNR{count}(j).y(1,:);
            allSimulations{count}(:,:,j)=sol_SNR{count}(j).y;
            
        end
        count = count+1;
    end
    tStimEnd_save=tStimEnd;
    save('./Results/stimTime2SigalAmplitude.mat','signalAmplitude','sol_SNR','allSimulations','tStimEnd_save');
    
else %else load the saved file
    load('./Results/stimTime2SigalAmplitude.mat');
    count = size(tStimEnd,2)+1;
end
simulation_UB = cellfun(@max,allSimulations,repmat({[]},1,count-1),num2cell(3*ones(1,count-1)),'UniformOutput',false); % get the maximum simulation value 
simulation_LB = cellfun(@min,allSimulations,repmat({[]},1,count-1),num2cell(3*ones(1,count-1)),'UniformOutput',false); % get the minimum simulation value 
%% find SNR

[rowIdx,colIdx] = find(min(signalAmplitude(:,[1:4],:),[],3)>2*mean(metData(1).Sigma_Y(:,[1:4])));
[~,tempIdx]=unique(colIdx);


%% plot max absolute amplitude vs stimulation time

f1=figure('units','normalized','outerposition',[0 0 1 1]);
titleText={'Lactate','Glutamate','Glucose','Aspartate','BOLD'};
% colour = linspecer(size(sol_SNR.y,2));

colour =[0.5  0.05 0.05
         0.95 0.53 0.13
         0    0.40 1.00
         0    0.40 0.20
         0.80 0    0
         0    0    1
         0.60 0    0.80
         1.00 0.12 0.36
         0.49 0.18 0.56
         0    0.60 0.87
         0    0.80 0.43
         0.69 0.35 0.73
         1.00 0.77 0.11
         1.00 0.70 0.58
         0.63 0.70 0.73
         0.64 0.47 0.11
         ];
     
AlphaLevel = 0.3; % set the alpha level for shaded areas in plots

ax1 = subplot(3,1,1);
hold on
timePoints2Plot_Idx = find(ismember(tStimEnd,[5 20 60 120 300 600 1200]));
xShift = [0.03 0.03 0.09 0.09 0.15 0.15 0.15 0.15];
% add lines at specific time points
for i = 1: size(timePoints2Plot_Idx,2)
   line(repmat(tStimEnd(timePoints2Plot_Idx(i)),2,1).',[0 55],'color',(colour(i,:)+(1-colour(i,:))*(1-AlphaLevel)),'LineWidth',5)% plota a line at each time pointe in the corresponding coulour
   text(tStimEnd(timePoints2Plot_Idx(i))-xShift(i),57.7,[num2str(tStimEnd(timePoints2Plot_Idx(i))),' s'],'FontWeight','bold','FontSize',32,'HorizontalAlignment','center'); %add the text lable at the time points
end

% plot max response curves
for i = 1:size(sol_SNR{1}(1).y,2)-2
    fill([tStimEnd.';flipud(tStimEnd.')],[min(signalAmplitude(:,i,:),[],3);flipud(max(signalAmplitude(:,i,:),[],3))],'b','FaceColor',(colour(end-1,:)+(1-colour(end-1,:))*(1-0.5)),'FaceAlpha',1)
    text(2e3,round(max(max(signalAmplitude(:,i,:),[],3)))+2,titleText{i},'FontSize',36,'FontWeight','bold'); %add the text label for each metabolite
end
% plot(tStimEnd,signalAmplitude(:,[1:4],1),'*','MarkerSize',10,'LineWidth',2);

% hleg = legend(titleText,'FontSize',32);
% hleg.AutoUpdate='Off'; %prevet Legend to auto add new lines
xlabel('Length of Stimulation [s]','FontSize',32)
ylabel(sprintf('Maximal signal amplitude\n relative baseline'),'FontSize',32)
set(ax1,{'XScale','XLim','FontSize','YTickLabels'},{'log',[0.5 1e4],32,strcat(ax1.YTickLabels,' %')});

maximizeAxes(f1,1,1);
setSubFigureNotations(ax1,32); %add the A-B notations to the subfigures
%% plot responses
f2=figure('units','normalized','outerposition',[0 0 1 1]);


count = 1;
for j = timePoints2Plot_Idx
    
    for i = 1:4
        ax(i) = subplot(3,2,i+2);
        hold on
        fill([timeVector;flipud(timeVector)],[simulation_LB{j}(:,i);flipud(simulation_UB{j}(:,i))],'b','FaceColor',colour(count,:),'FaceAlpha',AlphaLevel);

        title(titleText{i})
        ax(i).YLabel.String = 'Response (\Delta%)';
        ax(i).XLabel.String = 'Time [s]';
        set(ax(i),{'FontSize','XLim'},{32,[0 2000]});
    end

    count = count+1;
end
legend(ax(2),cellfun(@sprintf,repmat({'%d s'},1,size(timePoints2Plot_Idx,2)),num2cell(tStimEnd(timePoints2Plot_Idx)),'Uni',0),'Orientation','horizontal')
maximizeAxes(f2,2,2);
setSubFigureNotations(ax,32); %add the A-B notations to the subfigures
```
