## Supplementary Zip file for "Mechanistic model for human brain metabolism and its connection to the neurovascular coupling": README.md

# NVC_MetabolismModel 

Supplementary Material for paper named "Mechanistic model for human brain metabolism and the neurovascular coupling. "

To reproduce the results presented in the manuscript run the "Run.m" matlab script.
Please note that the "Generat models" require Matlab R2017b or earlier to run. 

TOOLBOXES
Please ensure that all of theses toolboxes are installed form their publically avilable sources and added to the matlab search path to ensure proper reproducabillity of the result. 
- The presented model implementation uses the “Advanced Multi-language Interface to CVODES and IDAS” (AMICI) toolbox (Fröhlich F et. al. Bioinformatics 2017) avilable from https://github.com/AMICI-dev/amici
- Parameter estimation alalysis were performed using the “Metaheuristics for systems biology and bioinformatics global optimization” (MEIGO) toolbox (Egea JA et. al., Bioinformatics, 2014) avilable from http://gingproc.iim.csic.es/meigom.html
- The MCMC-sampling analysis were performed using the "Parameter EStimation TOolbox" (PESTO) toolbox (Stapor P et. al. Bioinformatics 2018) avilable from https://github.com/ICB-DCM/PESTO


The folder titled "Matlab Files" contains all files necessary to reproduce the results presented in the paper. 
Here follows a shot decription of the most important of these files.

Folders: 
	Data - contains the data files used in a .mat-format.
	Models - contains the model files used for the various results.
	Parameters - Contains a .mat file with paramater values.
	Results - contains the results form the MCMC analysis and the stimulation time analysis.
Files:
	GenerateData.m 	- loads and structures the data files.
	GenerateModels.m- uses the AMICI amiwrap function to compile the .mex model files and the AMICI simulation functions.
	MCMCsampling.m 	- prefomres the MCMC analysis. 
	runParallelOpt.m- sets up and runs the runParameterEstimation_v2.m script used for parameter estimation. 

	simulateAndPlotMCMCresult.m - Plots the results from the MCMC analysis. Use this script to reproduce the main fiugres from the paper. 

The "Figures" folder contains all the figures in both .eps and .svg formats.
	
