## Supplementary Zip file for "Mechanistic model for human brain metabolism and its connection to the neurovascular coupling": Run.m

```
clear all
close all
clc

warning 'off'

addpath('..');


msg = "";
opts = ["Plot Figures" ....
        "Run simulations"...
        "Run parameter sampling"...
        "Run parameter estimation"...
        "Generate models"...
        ];

choice = menu(msg,opts);
addpath(genpath(fullfile(pwd,'Matlab Files')))
cd('./Matlab Files')
try

    switch choice
        case 1
            fprintf('Plotting Figures please wait ....')
           PlotResults
        case 2
            simulateAndPlotMCMCresult
        case 3
            MCMC_sampling
        case 4
            runParallelOpt
        case 5
            GenerateModels
    end
    
catch err
   close all 
   disp(err)
   fprintf('TOOLBOXES\nPlease ensure that all of theses toolboxes are installed form their publically avilable sources and added to the matlab search path to ensure proper reproducabillity of the result.\n- The presented model implementation uses the "Advanced Multi-language Interface to CVODES and IDAS" (AMICI) toolbox (Fröhlich F et. al. Bioinformatics 2017) avilable from https://github.com/AMICI-dev/amici\n- Parameter estimation alalysis were performed using the "Metaheuristics for systems biology and bioinformatics global optimization" (MEIGO) toolbox (Egea JA et. al., Bioinformatics, 2014) avilable from http://gingproc.iim.csic.es/meigom.html\n- The MCMC-sampling analysis were performed using the "Parameter EStimation TOolbox" (PESTO) toolbox (Stapor P et. al. Bioinformatics 2018) avilable from https://github.com/ICB-DCM/PESTO')
end
cd('../')
```
